# Highly-efficient Cpf1-mediated gene targeting in mice following high concentration pronuclear injection

**DOI:** 10.1101/097220

**Authors:** Dawn E. Watkins-Chow, Gaurav K. Varshney, Lisa J. Garrett, Zelin Chen, Erin A. Jimenez, Cecilia Rivas, Kevin S. Bishop, Raman Sood, Ursula L. Harper, William J. Pavan, Shawn M. Burgess

## Abstract

Cpf1 has emerged as an alternative to the Cas9 RNA-guided nuclease. Here we show that gene targeting rates in mice using Cpf1 can meet or even surpass Cas9 targeting rates (approaching 100% targeting) but require higher concentrations of mRNA and guide. We also demonstrate that co-injecting two guides with close targeting sites can result in synergistic genomic cutting, even if one of the guides has minimal cutting activity.

## INTRODUCTION

The CRISPR/Cas9 systems has revolutionized genome editing in many model and non-model organisms. Because of its high efficiency and simplicity of target design, CRISPR/Cas9 is now being used for a wide variety of molecular biology applications (Sander and Joung 2014). The technique was simplified with the use of single guide RNAs (sgRNA) containing a target site and chimeric crRNA and tracrRNA sequences that can direct Cas9 to the target site (Hsu *et al.* 2014). Target recognition only requires the presence of a protospacer adjacent motif (PAM) at the 3’ end of the target site. The widely used Cas9 protein from *Streptococcus pyogenes* (SpCas9) utilizes a G-rich, NGG PAM sequence that limits the selection of target sites, however, Cas9 orthologs and engineered Cas9 variants that recognize other PAM sequences have expanded the set of available target sites. Still more target sites are available with the recently identified CRISPR/Cpf1 class of proteins (Zetsche *et al.* 2015). Similar to Cas9 proteins, Cpf1 proteins are single RNA-guided endonucleases, but they function without tracrRNA and recognize a T-rich PAM. While an entire family of Cpf1 proteins was discovered, only two Cpf1 proteins, from *Acidaminococcus* (AsCpf1) and *Lachnospiraceae* (LbCpf1), have been shown to work for mammalian genome editing *in vitro*. Both AsCpf1 and LbCpf1 recognize a TTTN PAM site that could be useful in targeting regulatory regions or AT-rich genomes. Two groups have recently reported Cpf1 activity in mice, although with variable targeting rates and 3 different methods to deliver the Cpf1 into mouse zygotes including pronuclear microinjection of ribonuclear-protein complexes (RNPs), electroporation of RNPs, or cytoplamasmic microinjection of mRNA/gRNA mixtures (Hur *et al.* 2016; Kim *et al.* 2016). Here, we report the first demonstration of AsCpf1 activity using a distinct delivery method, pronuclear microinjection of RNA, show the targeting rate with this delivery method is highly dependent on RNA concentration, and that multiple tested target sequences and conditions were ineffective for targeting in zebrafish embryos.

## RESULTS AND DISCUSSION

To test if AsCpf1 had activity following pronuclear injection and if the targeting rate of AsCpf1 for *in vivo* gene knockout in mouse is affected by RNA concentration, Exon 1 of *Tyrosinase* (*Tyr*) was targeted with two different gRNAs (Figure 1A). The gRNAs were injected individually, or co-injected as a pair along with AsCpf1 RNA for pronuclear injection at either a “low” concentration (10ng/μl Cpf1 with 2.5ng each gRNA,) comparable to concentrations routinely used for Cas9 targeting, or a “high” concentration (50ng/μl with 100ng each gRNA). In total, 297 founders were screened following 13 injection sessions carried out in parallel to compare the targeting rate following injection of low or high concentrations of gRNA and Cpf1 RNA (Supplemental Table 1). Among the founder mice that were carried to term to allow for phenotype screening after birth, albino mice as well as mosaic mice with visible patches of albino hair were observed, phenotypes consistent with efficient targeting of the wild-type C57BL/6J *Tyr* allele in the hybrid C57BL/6J x FVB/N F1 mice (Figure 1B). The 297 founder mice were initially genotyped by sizing a fluorescently labeled PCR product spanning the two target sites to detect potential indels at any sites of DNA cleavage using both low resolution agarose gels and high resolution capillary electrophoresis (Supplemental Figure 1). In total, PCR products that differed from the expected 351bp wild-type product were detected in 73/297 founder mice, indicating that indels occurred with an overall frequency of 24.6%. Importantly, injection sessions with a high RNA concentration resulted in a much higher targeting rate (73.3% of the founders) compared to those with a low RNA concentration (3.4% of the founders) (Supplemental Table 1). Because of the overall high efficiencies of targeting with two guides at the high RNA concentration and small number of founders, we could not make conclusions about any effect the direction of the cross had on the targeting rate ((C57BL/6J female x FVB/N male) vs (FVB/N female x C57BL/6J male)) (Supplemental Table 2). However, in the low concentration injections, we observed a significantly higher targeting rate with C57BL/6J fertilized eggs (C57BL/6J female x FVB/N male) compared to the reciprocal cross (Supplemental Table 2). Importantly, the high concentration injections were more efficient independent of the direction of the cross (Supplemental Table 3).

**Figure 1.**
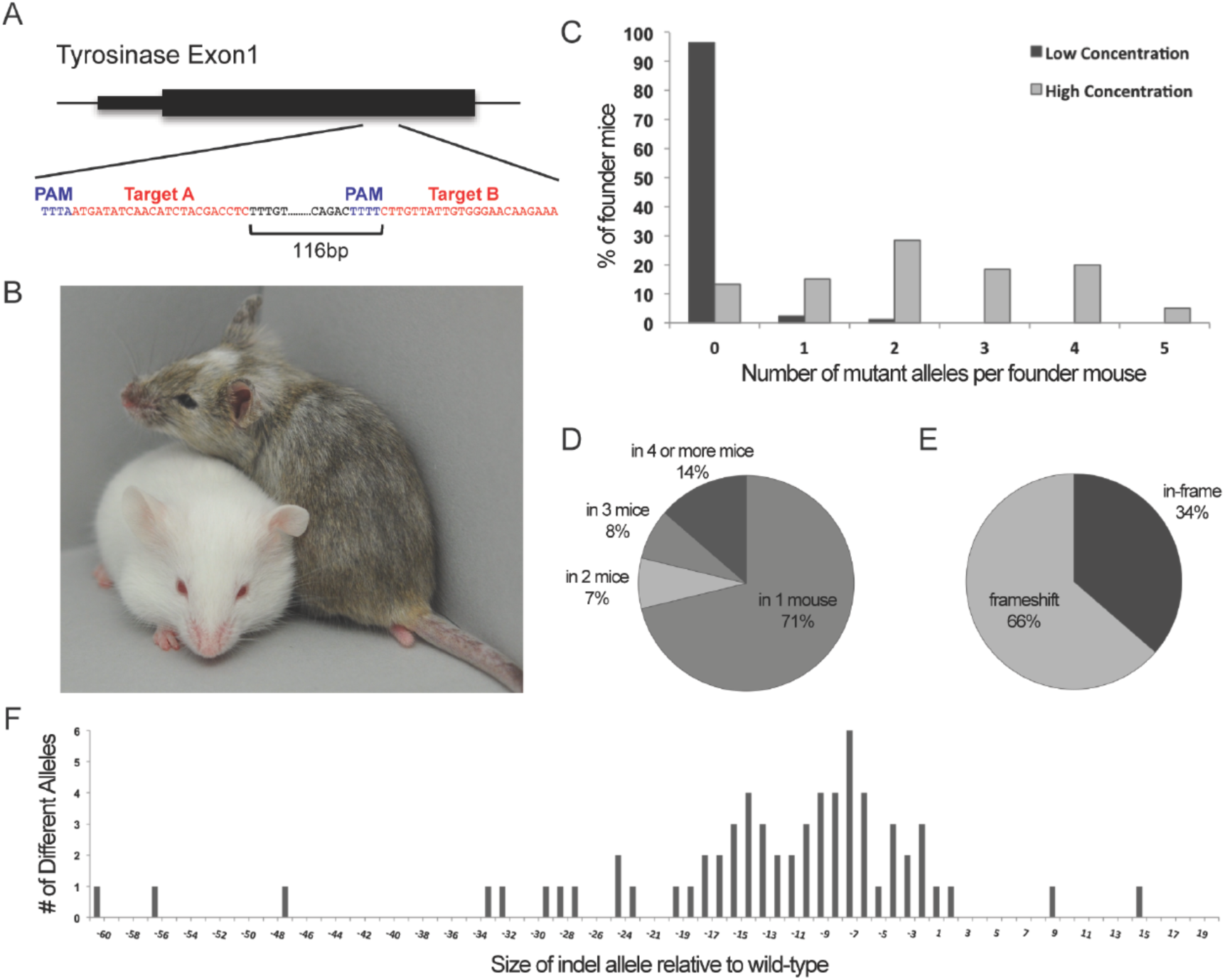
*In vivo* activity of AsCpf1. (A) Two gRNAs, A and B, were selected within *Tyrosinase* (*Tyr*) exon 1. (B) Two founder mice showing examples of a heterozygote albino mutant (left) next to a mosaic littermate (right), whose albino patches of fur resulted from AsCpf1-mediated mutation of the wild-type C57BL/6J *Tyr* allele within a subset of cells during development. (C) The percentage of founder mice carrying 0 to 5 different mutant *Tyr* alleles is shown for the low and high concentration injection sessions. (D) The percentage of mutant *Tyr* alleles that arose in a single founder or were recurring in two or more founder mice. (E) The percentage of mutant *Tyr* alleles predicted to introduce frameshift vs. in-frame mutations. (F) The net size of indel alleles compared to the expected size of the wild-type *Tyr* sequence.

To further characterize the AsCpf1-mediated indels, a combination of deep sequencing (Illumina MiSeq) and Sanger sequencing was carried out on PCR products spanning the target sites. This sequencing revealed that there was only one Target B site mutation; all other mutations were localized to the gRNA Target A site. Furthermore, in the more efficient high concentration injections, a higher targeting rate per 100 founders was observed at the Target A site when Target B was co-injected with 87.1% (54/62) of founders targeted following co-injection vs 42.8% (12/28) following single gRNA injected (Fisher’s test P<0.0001). While confirmation is needed from additional experiments, this intriguing result suggests that co-injection of multiple gRNAs may increase AsCpf1 activity.

In the low concentration injections, the majority of mice were wild-type (97%), and very few heterozygote or mosaic mice were generated that carried one (2%) or two (1%) mutant alleles. In contrast, the high concentration injections had both a higher rate of mutation and also a higher rate of mosaicism, with 5% of the founder mice each carrying as many as 5 different mutant alleles (Figure 1C). In total, 74 different alleles were identified in the 52 mutant founder mice that were fully characterized by deep sequencing. The majority of these alleles (71%) arose only in a single founder, however some alleles (29%) were reoccurring, as 9 alleles arose independently in 4 or more founder mice (Figure 1D). As expected, approximately 2/3 of the alleles introduced a frameshift (Figure 1E). The indels ranged in size from −60bp to +15bp and were mostly deletion alleles (Figure 1F and Supplemental Figure 2), consistent with observations in *Drosophila melanogaster* (Port and Bullock 2016). All of the founder mice were confirmed to be heterozygous for a SNP (rs31191169) known to be polymorphic between C57BL/6J and FVB/N that is located 195bp proximal to the Target A site, indicating that no large deletions occurred that extended beyond the PCR amplicon used for sequencing.

The majority of the founder mice appeared mosaic, as indicated by an uneven ratio of alleles within the PCR product where one or more mutant alleles were detected at less than the expected 50%. This result is consistent with the mosaic coat color pattern observed in founder mice carried to term and indicates that DNA cleavage often occurred after the one cell stage. Because of this high frequency of mosaic founders, germline transmission of the AsCpf1-induced mutant alleles was confirmed by breeding 6 founder mice to FVB/N. One of these founders was albino and 5 were mosaic with a range of 10-70% estimated albino coat color contribution, representing a wide range of the observed mosaicism including the lowest and highest contribution founders (Supplemental Figure 3). In all 6 mice, the Cpf1-mediated mutant allele was transmitted and in one mosaic mouse, two different mutant alleles were transmitted.

In contrast to mouse, we were unable to successfully induce gene knockout in zebrafish with AsCpf1. Using injection methods previously successful with Cas9 (Varshney *et al.* 2015), we targeted 6 genes with 40 different targets (23nt or 24nt), but no activity was detected by CRISPR-STAT (Carrington *et al.* 2015). These results suggest that either AsCpf1 has lower activity in zebrafish than Cas9 or that there are unique technical requirements for AsCpf1 activity in this species.

In summary, we detected AsCpf1 activity in mouse, but not in zebrafish. The level of AsCpf1 activity in mouse zygotes was highly dependent on RNA concentration and at high concentrations was extremely effective, confirming that AsCpf1 provides an excellent alternative to Cas9 for the generation of knockout mice. Our targeting rate with AsCpf1 at high RNA concentrations (42-100% of founders) is significantly higher than at low RNA concentrations (0-21%) and is consistent with other studies that used a variety of delivery methods for Cpf1-medicated targeting in mice. For comparison, successful AsCpf1 targeting was observed following pronuclear microinjection of RNPs (17-83%) (Hur *et al.* 2016), electroporation of RNPs (14-64%) (Hur *et al.* 2016), and cytoplasmic microinjection of RNA (18-79%) (Kim *et al.* 2016) (Supplemental Table 4).

Our “high” concentration is comparable to that published for cytoplasmic microinjection of RNA (Kim *et al.* 2016), however much higher than required for Cas9 activity with pronuclear microinjections which typically are performed using lower RNA concentrations than cytoplasmic microinjections (Singh *et al.* 2015). Because of the different delivery methods and limited number of guides tested, few conclusions can be drawn about the most important variables affecting the AsCpf1 targeting rate in mice, which may include RNA concentration, mouse strain, delivery method, or local target sequence. Importantly, our study provides the first direct comparison of a guide RNA at different concentrations and suggests that Cpf1 is ineffective for targeting at an RNA concentration that is effective for Cas9 in our laboratory and others (Mashiko *et al.* 2013, Yang *et al.* 2013, Horii *et al.* 2014). Interestingly, we observed that targeting was more efficient with co-injection of two adjacent guides, even though one guide was essentially inactive. While this represents data from only a single locus, the statistically significance (P<0.0001) indicates that the effect of co-injection on targeting rate is worth testing more broadly in the future.

Because the T-rich PAM sequences recognized by AsCpf1 differ from Cas9, the combination of these nucleases will permit the precise targeting necessary for knock-in projects, especially those aimed at modeling short sequence variants at particular genomic locations. Also, the shorter size of the gRNA utilized by AsCpf1 provides an advantage over Cas9 for reducing the cost of RNA oligonucleotide synthesis and has the potential to improve efficiency of gRNA delivery. Importantly, the frequency of homology-directed repair or ligation-mediated insertion by Cpf1 remains to be determined and could yet provide the largest advantage for Cpf1 over Cas9.

## METHODS

### Preparation of AsCpf1 RNA and gRNA

AsCpf1 targets were identified with Benchling software and selected based only on the presence of a predicted PAM site without considerations of predicted efficiency. gRNAs were synthesized as custom RNA oligonucleotides with standard desalting (Integrated DNA Technologies), and resuspended in Picopure system purified water (Hydro Systems) at 500 or 1,000ng/μl. Two different gRNAs were synthesized for mouse pronuclear injections (unique target sequence for each is indicated in bold): Target A(5’TAATTTCTACTCTTGTAGAT**ATGATATCAACATCTACGACCTC**) and Target B (5’TAATTTCTACTCTTGTAGAT**CTTGTTATTGTGGGAACAAGAAA**). The hAsCpf1 plasmid (Addgene 69982) was a gift from Feng Zhang. A PCR product amplified from the hAsCpf1 plasmid (Cpf1_Fwd actggcttatcgaaattaatacgactc; Cpf1_Rev ccccagctggttctttcc) was used as a template for *in vitro* RNA synthesis using mMessage mMachine T7 Ultra Transcription Kit (Thermo Fisher/Ambion) according to the manufacturer’s instructions. RNA was recovered using the MEGAclear Transcription Clean-up Kit (Thermo Fisher/Ambion). RNA quality was assessed with a Bioanalyzer instrument (Agilent Genomics) and stored in aliquots at −80°C until used for injections

### Pronuclear Injection

Pronuclear injection was performed with standard procedures (Behringer *et al.* 2014). Briefly, fertilized eggs were collected from superovulated FVB/N (Taconic Farms; abbreviated FVB) or C57BL/6J (Jackson Laboratories; abbreviated B6) females approximately 9 hours after mating with C57BL/6J or FVB/N males, respectively. Microinjections were performed using a capillary needle with a 1-2um opening pulled with a Sutter P-1000 micropipette puller. The pronucleus was injected using a FemtoJet 4i (Eppendorf) with continuous flow that we estimate to result in approximately 2pl of injection mix. Following visualization of pronuclear swelling, the needle was pulled out through the cytoplasm, likely resulting in a small amount of additional RNA delivery to the cytoplasm. Injected eggs were surgically transferred to pseudopregnant CB6F1 hybrid recipient females, bred at the NIH from a cross of Balb/cJ females to C57Bl/6J males. In general, the injection mix contained AsCpf1 RNA and gRNA diluted in 10mM Tris, 0.25mM EDTA (pH 7.5). Specific RNA and DNA concentrations for each injection session are provided (Supplemental Table 1).

### Mouse husbandry

All animal procedures were performed in a pathogen-free, AAALAC-approved facility in accordance with NIH guidelines and approved by the NHGRI Animal Care and Use Committee (ACUC).

### Genotyping

Founder animals were screened using a combination of PCR and sequencing as previously described (Varshney *et al.* 2015). Briefly, DNA was extracted from midgestation whole embryos or pup tail biopsies and purified using a Gentra Puregene Mouse Tail Kit (Qiagen). A PCR product spanning both gRNA target sites was generated using a Tyr-specific forward primer with an M13-tail (tgtaaacgacggccagtTCTATGTCATCCCCACAGGCAC) and a *Tyr*-specific reverse primer containing a pig-tail (gtgtcttGGTGACGACCTCCCAAGTACTC). The PCR product was amplified with the addition of a 6-FAM or HEX labeled M13-forward oligonucleotide and run on an ABI 3130xl with ROX400 or ROX500 size standards to detect small indels (<50bp) at single base pair resolution. Standard agarose gels were also used to screen PCR products for larger indels (>20bp) including possible deletion of the 116bp between the two guides which was never observed.

A subset of founders with PCR product sizes that differed from wild-type were further characterized by Sanger sequencing of total PCR products or subcloned PCR products generated with additional gene specific primers. Additionally, selected founders were characterized with deep sequencing using PCR products generated with *Tyr*-specific primers (tgtaaaacgacggccagtTTTTCTTACCTCACTTTAGCAAAACA and GGATGCTGGGCTGAGTAAGT) along with the barcoded third primer set, as described earlier (Varshney *et al.* 2015). The sequences were analyzed using minor modifications of the amplicoDIVider pipeline initially developed to identify deletion and insertion variants (DIVs) in deep sequencing data from zebrafish Cas9 mutants (Varshney *et al.* 2015). The amplicoDIVider scripts are publicly available (https://research.nhgri.nih.gov/software/amplicondivider).

## ACKNOWLEDGEMENTS

We thank Feng Zhang and Bernd Zetsche for providing the AsCpf1 plasmid and valuable advice, Arturo Incao for genomic DNA preparation and PCR amplification, Gene Elliott for assistance with microinjections, Laura Baxter for careful reading of the manuscript, and Abdel Elkahloun and Weiwei Wu of the NHGRI Microarray Core for assistance assessing RNA quality. This research was supported by the Intramural Research Program of the National Human Genome Research Institute at the National Institutes of Health (SMB: ZIA HG000183-15 and WJP: HG000136-11)

---

**Supplemental Figure 1.**
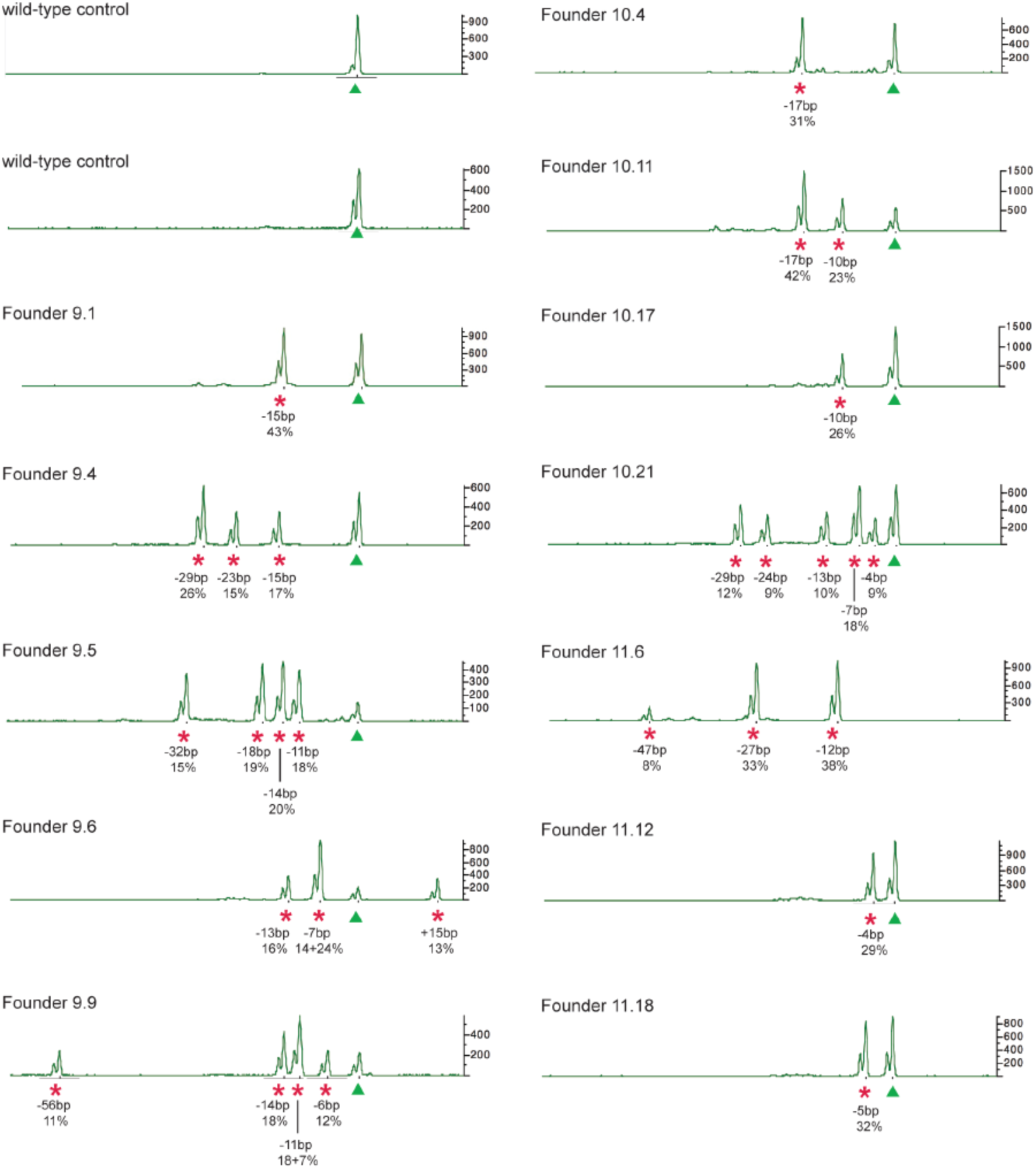
Examples of 2 wild-type and 12 founder mice screened with a fluorescently labeled PCR product. WT *Tyr* allele products are indicated (green arrowhead) as well as mutant *Tyr* alleles confirmed by deep sequencing of an independent PCR product (red *). For mutant alleles, the size of each allele relative to wild-type is indicated along with the percentage of sequencing reads. Note that founder 11.6 lacks a wild-type allele. Upon sequencing, we determined that the single −7bp peak in founder 9.6 was actually a mix of two different −7bp mutant alleles. Similarly the −11bp peak in founder 9.9 contained two different −11bp mutant alleles.

**Supplemental Figure 2.**
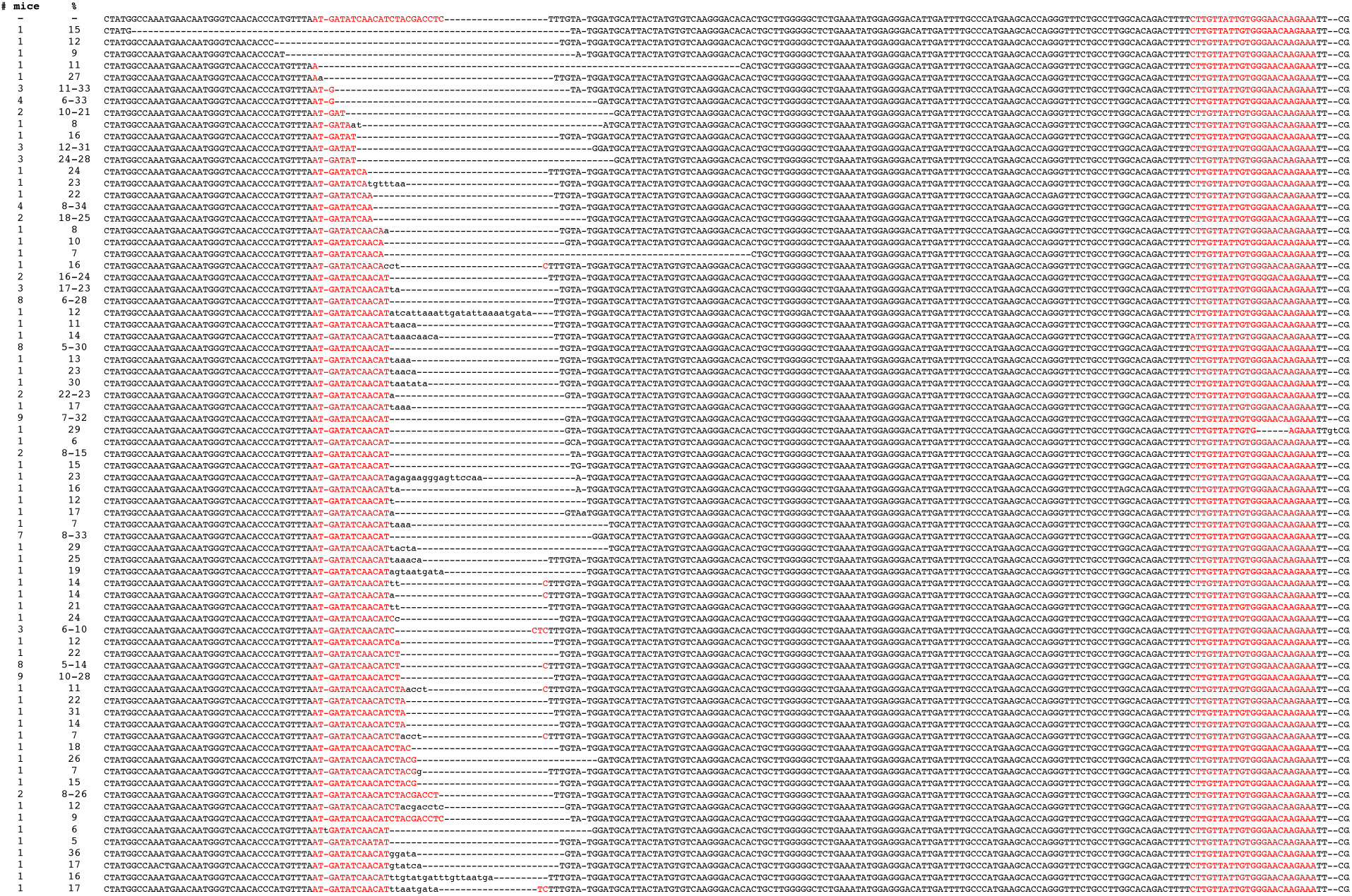
Sequence alignment for 74 alleles compared to the expected wild-type *Tyrosinase* sequence. The target site is indicated (red) along with deleted bases (–) and inserted or altered bases (lower case). Note that in some cases the exact breakpoint cannot be determined from the sequence data because bases on either side of the breakpoint are identical. For each allele, the number of founder mice observed carrying that allele is indicated along with the percentage of sequencing reads within the founder PCR product. For alleles observed in multiple founders, a range of percentages is given.

**Supplemental Figure 3.**
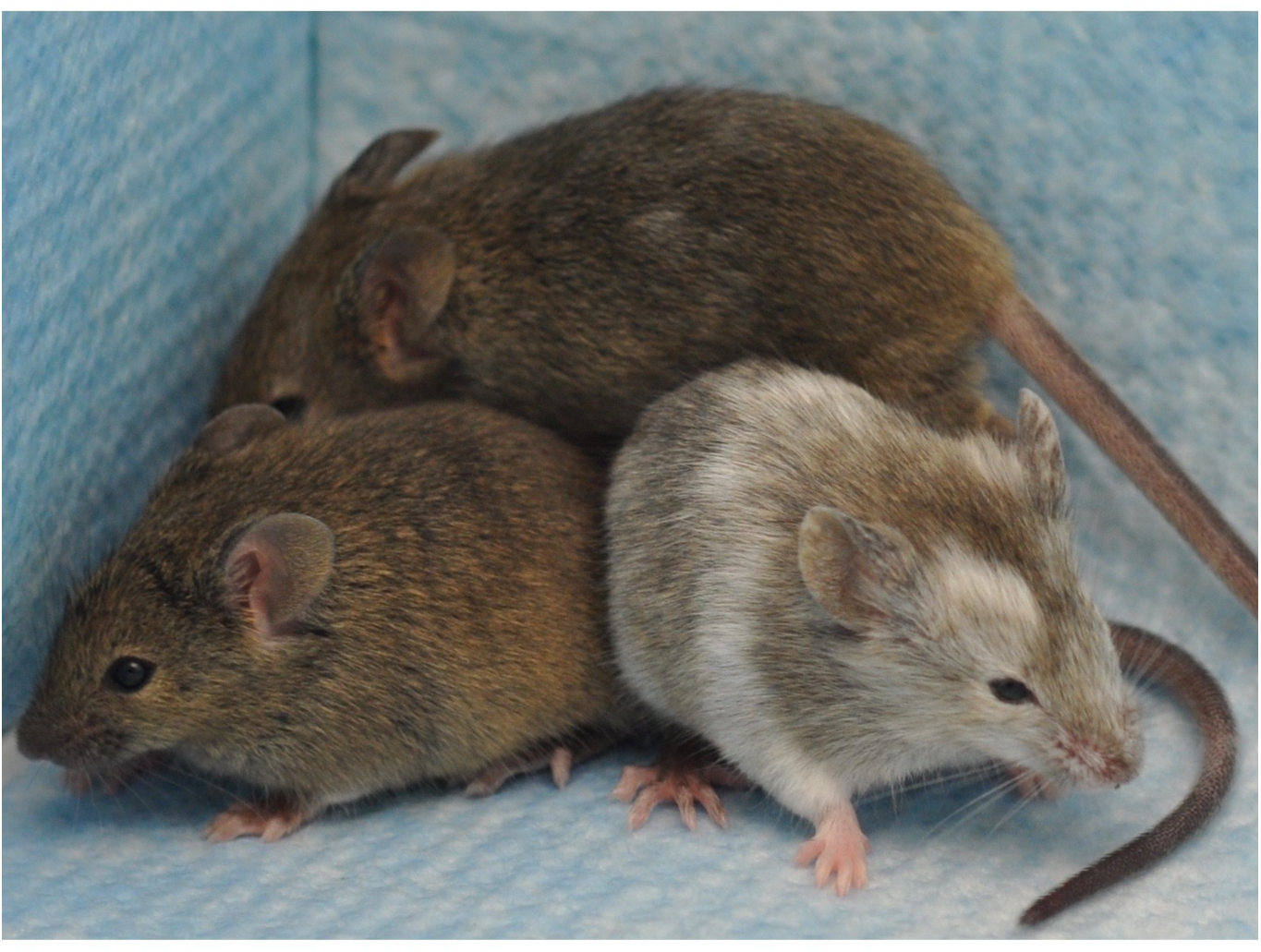
Mosaic transmission of AsCpf1-mediated mutant alleles. 6 founders were bred to confirm transmission of the mutant alleles: 1 albino mouse and 5 mosaics ranging from 10-70% albino coat color contribution. In crosses of all 6 mice to FVB, germline transmission of the mutant allele occurred. Two of the 6 mice are shown here: the mouse with the lowest albino coat color contribution (10%) in the center, and the mouse with the highest coat color contribution (70%) on the right. For comparison, a wild-type agouti mouse is shown on the left.

**Supplemental Table 1.**
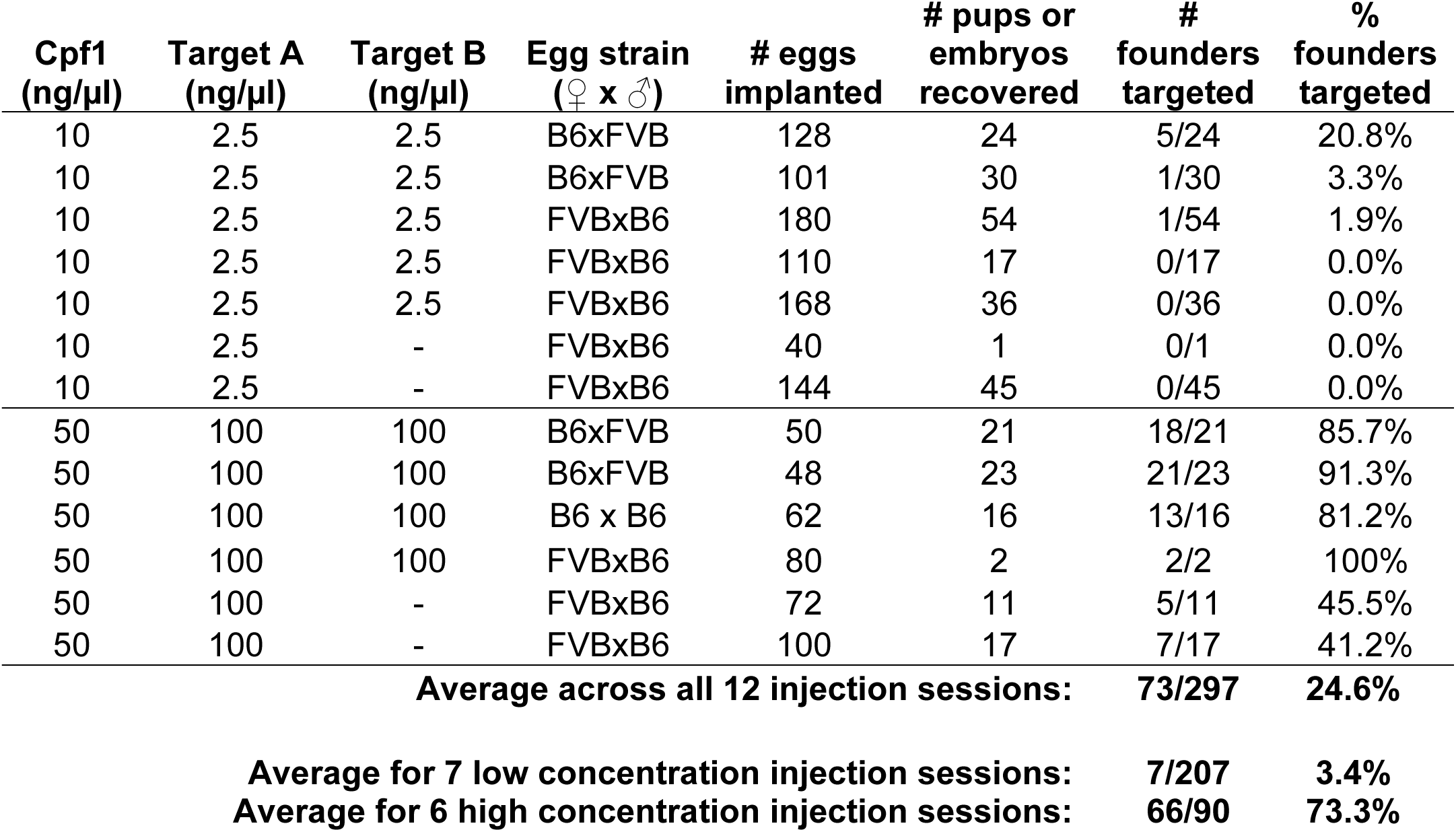
Summary of injection sessions

| **Cpf1**<br>**(ng/μl)** | **Target A**<br>**(ng/μl)** | **Target B**<br>**(ng/μl)** | **Egg strain**<br>**(♀ x ♂)** | **# eggs**<br>**implanted** | **# pups or**<br>**embryos**<br>**recovered** | **#**<br>**founders**<br>**targeted** | **%**<br>**founders**<br>**targeted** |
| --- | --- | --- | --- | --- | --- | --- | --- |
| 10 | 2.5 | 2.5 | B6xFVB | 128 | 24 | 5/24 | 20.8% |
| 10 | 2.5 | 2.5 | B6xFVB | 101 | 30 | 1/30 | 3.3% |
| 10 | 2.5 | 2.5 | FVBxB6 | 180 | 54 | 1/54 | 1.9% |
| 10 | 2.5 | 2.5 | FVBxB6 | 110 | 17 | 0/17 | 0.0% |
| 10 | 2.5 | 2.5 | FVBxB6 | 168 | 36 | 0/36 | 0.0% |
| 10 | 2.5 | - | FVBxB6 | 40 | 1 | 0/1 | 0.0% |
| 10 | 2.5 | - | FVBxB6 | 144 | 45 | 0/45 | 0.0% |
| 50 | 100 | 100 | B6xFVB | 50 | 21 | 18/21 | 85.7% |
| 50 | 100 | 100 | B6xFVB | 48 | 23 | 21/23 | 91.3% |
| 50 | 100 | 100 | B6 x B6 | 62 | 16 | 13/16 | 81.2% |
| 50 | 100 | 100 | FVBxB6 | 80 | 2 | 2/2 | 100% |
| 50 | 100 | - | FVBxB6 | 72 | 11 | 5/11 | 45.5% |
| 50 | 100 | - | FVBxB6 | 100 | 17 | 7/17 | 41.2% |
| **Average across all 12 injection sessions:** |  |  |  |  |  | **73/297** | **24.6%** |
| **Average for 7 low concentration injection sessions:** |  |  |  |  |  | **7/207** | **3.4%** |
| **Average for 6 high concentration injection sessions:** |  |  |  |  |  | **66/90** | **73.3%** |

**Supplemental Table 2.** Effect of strain on targeting rate for each injection concentration*

|  | **B6 ♀ x FVB ♂** | **FVB ♀ x B6 ♂** | **Comparison**<br>**(Fisher's)** |
| --- | --- | --- | --- |
| **Total for all injection sessions** | 45/98 (45.9%) | 15/183 (8.2%) | **P < 0.0001** |
| **High concentration with 2 guides** | 39/44 (88.6%) | 2/2 (100%) | **P = 1.0000** |
| **High concentration with 1 guide** | - | 12/28 (42.9%) | - |
| **Low concentration with 2 guides** | 6/54 (11.1%) | 1/107 (0.9%) | **P = 0.0060** |
| **Low concentration with 1 guide** | - | 0/46 (0%) | - |

**Supplemental Table 3.** Effect of concentration on targeting rate for each strain *

|  | **High**<br>**Concentration** | **Low**<br>**Concentration** | **Comparison**<br>**(Fisher's)** |
| --- | --- | --- | --- |
| **B6 ♀ x FVB ♂, all injections** | 39/44 (88.6%) | 6/54 (11.1%) | **P < 0.0001** |
| **B6 ♀ x FVB ♂ with 2 guides** | 39/44 (88.6%) | 6/54 (11.1%) | **P < 0.0001** |
| **B6 ♀ x FVB ♂ with 1 guide** | - | - | - |
| **FVB ♀ x B6 ♂, all injections** | 14/30 (46.7%) | 1/153 (0.7%) | **P < 0.0001** |
| **FVB ♀ x B6 ♂ with 2 guides** | 2/2 (100%) | 1/107 (0.9%) | **P = 0.0005** |
| **FVB ♀ x B6 ♂ with 1 guide** | 12/28 (42.9%) | 0/46 (0%) | **P < 0.0001** |

**\*Data in this table is tallied from Supplemental Table 1**

**Supplemental Table 4:**
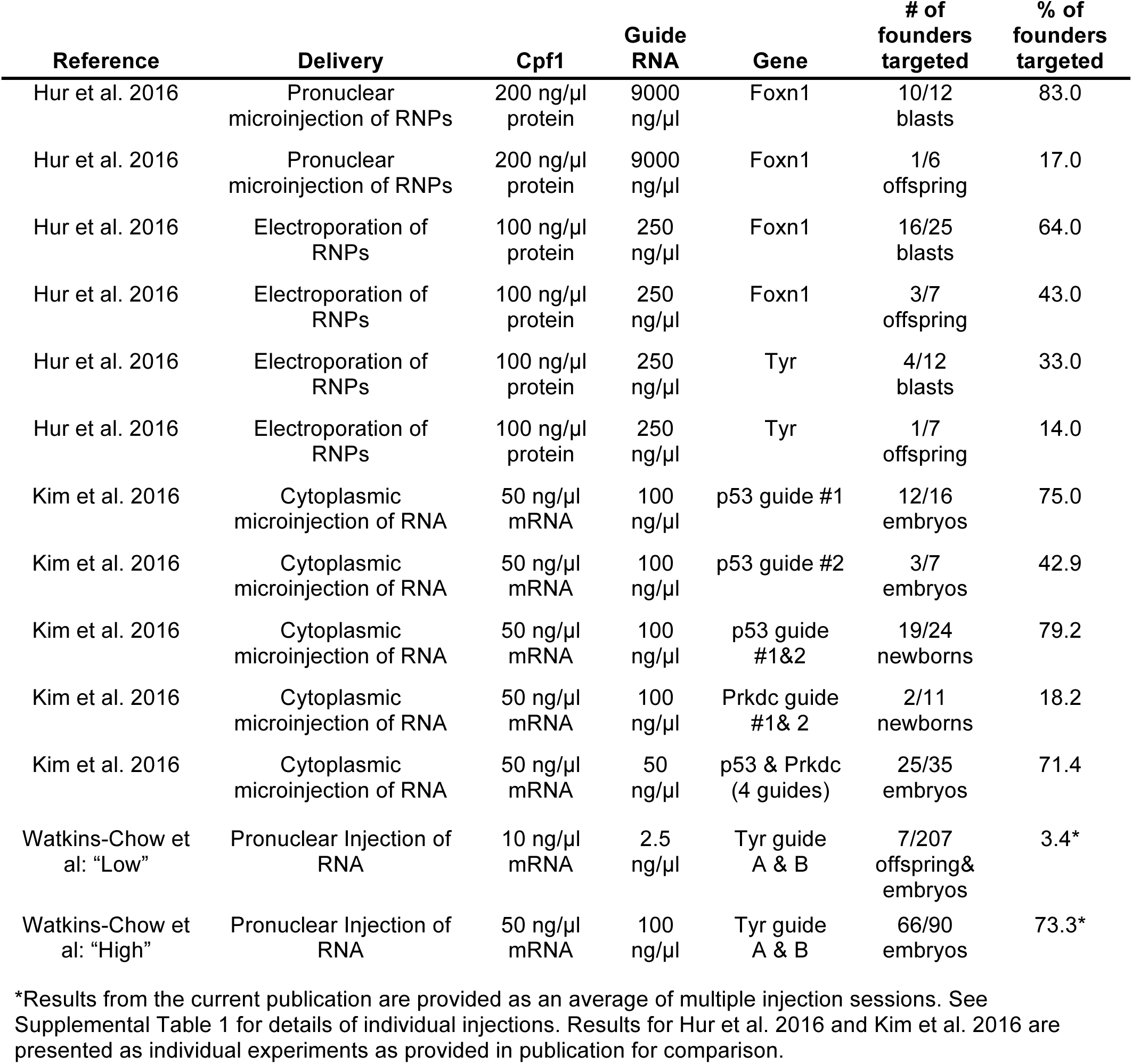
Comparison of successful targeting with AsCpf1

| Reference | Delivery | Cpf1 | Guide RNA | Gene | # of founders targeted | % of founders targeted |
| --- | --- | --- | --- | --- | --- | --- |
| Hur et al. 2016 | Pronuclear microinjection of RNPs | 200 ng/μl protein | 9000 ng/μl | Foxn1 | 10/12 blasts | 83.0 |
| Hur et al. 2016 | Pronuclear microinjection of RNPs | 200 ng/μl protein | 9000 ng/μl | Foxn1 | 1/6 offspring | 17.0 |
| Hur et al. 2016 | Electroporation of RNPs | 100 ng/μl protein | 250 ng/μl | Foxn1 | 16/25 blasts | 64.0 |
| Hur et al. 2016 | Electroporation of RNPs | 100 ng/μl protein | 250 ng/μl | Foxn1 | 3/7 offspring | 43.0 |
| Hur et al. 2016 | Electroporation of RNPs | 100 ng/μl protein | 250 ng/μl | Tyr | 4/12 blasts | 33.0 |
| Hur et al. 2016 | Electroporation of RNPs | 100 ng/μl protein | 250 ng/μl | Tyr | 1/7 offspring | 14.0 |
| Kim et al. 2016 | Cytoplasmic microinjection of RNA | 50 ng/μl mRNA | 100 ng/μl | p53 guide #1 | 12/16 embryos | 75.0 |
| Kim et al. 2016 | Cytoplasmic microinjection of RNA | 50 ng/μl mRNA | 100 ng/μl | p53 guide #2 | 3/7 embryos | 42.9 |
| Kim et al. 2016 | Cytoplasmic microinjection of RNA | 50 ng/μl mRNA | 100 ng/μl | p53 guide #1&2 | 19/24 newborns | 79.2 |
| Kim et al. 2016 | Cytoplasmic microinjection of RNA | 50 ng/μl mRNA | 100 ng/μl | Prkdc guide #1& 2 | 2/11 newborns | 18.2 |
| Kim et al. 2016 | Cytoplasmic microinjection of RNA | 50 ng/μl mRNA | 50 ng/μl | p53 & Prkdc (4 guides) | 25/35 embryos | 71.4 |
| Watkins-Chow et al: "Low" | Pronuclear Injection of RNA | 10 ng/μl mRNA | 2.5 ng/μl | Tyr guide A & B | 7/207 offspring& embryos | 3.4* |
| Watkins-Chow et al: "High" | Pronuclear Injection of RNA | 50 ng/μl mRNA | 100 ng/μl | Tyr guide A & B | 66/90 embryos | 73.3* |

\*Results from the current publication are provided as an average of multiple injection sessions. See Supplemental Table 1 for details of individual injections. Results for Hur et al. 2016 and Kim et al. 2016 are presented as individual experiments as provided in publication for comparison.

